# The applying of immuno-RCA for the high-sensitivity detection of the ABO blood group antibodies on the printed glycoarray

**DOI:** 10.1101/2024.12.11.625986

**Authors:** E.E. Kornilova, R.R. Kutukov, S.M. Polyakova, A.Yu. Nokel, S.K. Zavriev, D.Yu. Ryazantsev

## Abstract

Detecting small amounts of analytes, especially antibodies, presents a significant challenge in high-throughput methods. Fortunately, nucleic acid amplification techniques provide a promising solution. We have successfully developed and tested an innovative technology for detecting anti-glycan antibodies, utilizing a printed glycan array combined with a rolling circle amplification reaction based on the ABO blood group antibody model. This breakthrough has dramatically enhanced the sensitivity of our immunoassay, improving it by over an order of magnitude and allowing us to detect concentrations as low as 1 ng/ml. This advancement opens new approaches for research and clinical applications, making previously undetectable analytes accessible for study.

## Introduction

The accurate detection and quantification of immunoglobulins in different human biological liquids (blood serum, ascites etc.) and tissues play a critical role in the diagnosis and monitoring of numerous diseases, including cancer. Recently, monoclonal and polyclonal anti-glycan antibodies (AGA) against bacterial polysaccharides during infectious deceases, tumor-associated carbohydrate antigens (TACA) in cancer, glycans of carbohydrate-related blood group antigens in blood transfusions and others have become increasingly important in this field (Temme et al., 2021).

In recent years, advancements in molecular biology and biotechnology have introduced highly sensitive methods to detect antibodies, even at low concentrations. Among these innovations, using rolling circle amplification (RCA), particularly in its immunological application (Immuno-RCA), has emerged as a powerful technique for enhancing detection sensitivity (Zhang et al., 2024, Zhou et al., 2004). Immuno-RCA combines the specificity of antibody-antigen interactions with the amplification capability of RCA, significantly improving the detection limit for various antibodies. This method has demonstrated advantages over traditional techniques like enzyme-linked immunosorbent assays (ELISAs), especially when dealing with low-abundance proteins or antibodies.

The application of Immuno-RCA has proven especially valuable in assays, where the detection of multiple immunoglobulins (such as IgG, IgA, and IgM) is required (Horta et al., 2021). In cancer research, this method allows for the profiling of antibodies that bind to TACAs, aiding in identifying tumor cells. Moreover, the ability of Immuno-RCA to provide high sensitivity and specificity in detecting even trace amounts of immunoglobulins opens new possibilities for early cancer detection and targeted therapies.

In this study, we explore the potential of Immuno-RCA in the high-sensitive detection of anti-glycan immunoglobulins against the blood group ABO antigens printed on a microscope size slide (microchip).

## Material and Methods

### Biological samples

Group A blood sample was collected from healthy adult donor. Donor has signed an informed consent form approved by the local ethics committee of A.V. Vishnevsky Institute of Surgery (2024, Moscow, Russia). One hour after blood collection, sample was centrifuged for 10 min at 1500×g. Serum was stored at 80°C until analysis.

Microchips. Aminospacered A type 2 and B type 2 (GlycoNZ, New Zealand) were printed onto N-hydroxysuccinimide-activated slides (Semiotik LLC, Russia) at 100 μM concentrations in 300 mM phosphate buffer solution containing 0.001% Tween20 (SigmaAldrich, USA) and 50% relative humidity in a non-contact manner using SciFlexArrayer S5, Scienion (drop size about 0.9 nl). The ligands were combined into eight blocks which contained four repeats of each ligand at 100 μM concentrations. The Cy3 labeled BSA (0.025-0.5 μg/mL) and streptavidin were also printed as a control spots. The slide scheme in depth is shown in the Fig.S1.

Antibodies and Proteins. To detect anti-human immunoglobulins, biotinylated goat anti-human IgG, IgM, and IgA (H+L) antibody (A18851, Thermo Fisher Scientific) were used. The fluorescent-labeled BSA was obtained by the reaction between the BSA (Sisco Research Laboratories) and sulfo-Cy3 N-hydroxysuccinimideester (Lumiprobe) using the manufacturer protocol. Alexa Fluor™ 555 labeled streptavidin (S32355) was from Thermo Fisher Scientific, USA, non-labeled streptavidin (nrpa05s-1k) was from Nuptec, China. The phage Phi29 DNA polymerase was expressed in the E. coli strain M15 and purified in the laboratory of molecular diagostics IBCH RAS. The detailed protocol is described in the supplementary.

Oligonucleotides. Oligonucleotides used in this work are shown in Table S1.

Circular template preparation. Circular templates were obtained as described in Goryunova et. al., 2021 using oligonucleotides #2 and ##3-9 (Table S1). The circular template concentration was determined using QuDye ssDNA Assay Kit (Lumiprobe) and linear template oligonucleotides as concentration standards.

RCA kinetic assay. The polymerization kinetic test was carried out to estimate the rolling circle amplification reaction rate and phi29 DNA polymerase activity. Reaction mix (20 μL) contained RCA buffer (33 mM Tris-acetate pH 8, 10 mM magnesium acetate, 66 mM potassium acetate, 0.1% Tween 20, 1 mM DT), 0.83 mM of dATP, dTTP and dGTP, 1 μM of primer pr25WoCtBio, 1 μM of circular template, 5 u of phi29 DNA polymerase, 0.5 μM of fluorescent-labeled probe beacWoC3. Different amounts of hand-made phi29DNA polymerase was used in contrast to Thermo-Fisher Scientific phi29 DNA polymerase (EP0091) to measure the activity of our enzyme. RCA reaction was carried out in DT96 PCR machine at 30C for 3 h, the fluorescence was detected every 15 secs in the HEX channel. After the assay was complete the raw data were collected and scatter plots for each sample were plotted. The initial linear parts of the plots were determined and linear fitting was performed for them. The slope of the line equation (y = mx + b, where the slope is equal to m) represents the amplification rate.

Conventional immunoassay. All the incubations were carried out at room temperature and weak mixing in a humid chamber (~30 rpm). Microchip was blocked with 150 mM borate buffer containing 50 mM ethanolamine (SigmaAldrich, USA), pH 8.5 for two hours at room temperature and shaking (100 rpm). Then printed area was restricted by a hydrophobic marker (PapPen, USA) and chip was placed in a humid chamber (~70% relative humidity). The sample diluted in the buffer TBS containing 1% BSA and 0.1% of Tween20 (standard incubation buffer) was applied (0.7 ml) on the microchip surface and incubated for 1 h at 70% relative humidity and slow shaking (30 rpm), then washed in TBS with 0.05% Tween 20 three times (standard washing). 700 μL of 1 ug/ml goat anti-human biotinylated antibody diluted in standard incubation buffer was applied and the microchip was incubated for 1 h as above. The standard washing was performed and 700 μL 0.5 μg/mL streptavidin-Alexa Fluor™ 555 conjugate diluted in standard incubation buffer was applied on the chip and incubated for 1 h as above. The standard washing was performed followed by washing with deionized water and spin drying. Microchip was scanned using InnoScan 1100 (Innopsys, France) at 10 μm resolution, the obtained images were analyzed in the ScanArray Express 4.0 software (Perkin Elmer, USA).

Immuno-RCA. The initial steps were identical to the same in the previous paragraph. After incubation with anti-human antibody and washing the non-labeled streptavidin solution (0.5 ug/ml in standard incubation buffer) was applied to microchip. The complex of the circular template and biotinylated primer (1 nM in standard incubation buffer supplemented with 50 ng/μL yeast RNA, SigmaAldrich) was prepared by annealing biotinylated primer pr25WoCtBio and circular template. This complex was applied to the chip surface after standard washing and microchip was incubated for 1 hour. The RCA reaction mix was prepared (1 x RCA buffer, 0.75 mM 0.83 mM of dATP, dTTP, and dGTP, 0.5 μM probe beacWoC3, 0.5 u phage phi29 DNA polymerase, 0.2% of BSA, 200 ng/μL of yeast RNA) and applied to the microchip after standard washing. Rolling circle amplification was carried out during 1-3 hours at room temperature. The standard washing was performed and the microchip was washed with deionized water. Water was removed completely and microchip was scanned using InnoScan 1100 (Innopsys, France) with a resolution of 10 μm, the obtained images were analyzed in the ScanArray Express 4.0 software (Perkin Elmer, USA). The mean signal intensities (expressed in relative fluorescent units, RFU) within replicates with standard deviation were calculated. The signal was considered significant if it exceeded 10 average fluorescence values of the chip surface without spots (1500 RFU).

## Results and discussion

### Phi29 phage polymerase expression and purification, activity determination

We obtained about 10 mg of pure phi29 phage DNA polymerase per 1 L of bacterial culture. The enzyme was at least 95% pure (see Fig S1). In a comparison of phi29 DNA polymerase from Thermo Fisher, the activity test has shown that protein concentration equal 0.3 mg/ml corresponds to activity equal to 10 u/μL.

### Choice of template length and RCA reaction conditions

The optimal reaction conditions that were established and described in the method section. Under those conditions, we obtained the highest fluorescence increase during RCA.

To choose the optimal template length we carried out the RCA polymerization reaction kinetics test with different templates using the method described in the Material and Methods section. The lengths of tested circular templates were 55-64 nucleotides. Only three deoxynucleotide triphosphates (dATP, dGTP, and dTTP) were used to decrease non-specific amplification according to (Zingg JM, Daunert S, 2018) so we designed our templates without G-base. We observed a smooth increase in the amplification rate from 55 nucleotide long template to 58 nucleotide long template and a smooth decrease in the reaction rate from 58 to 64 nucleotide long template (Fig.1). These data are partially consistent with Joffroy et al 2017 showed that RCA demonstrated a sinusoidal template length-dependent amplification rate and the highest RCA rate (with steps between maximum equal ~10.3 pair) observed for 69 pairs long circular template and decreases with length increasing. The

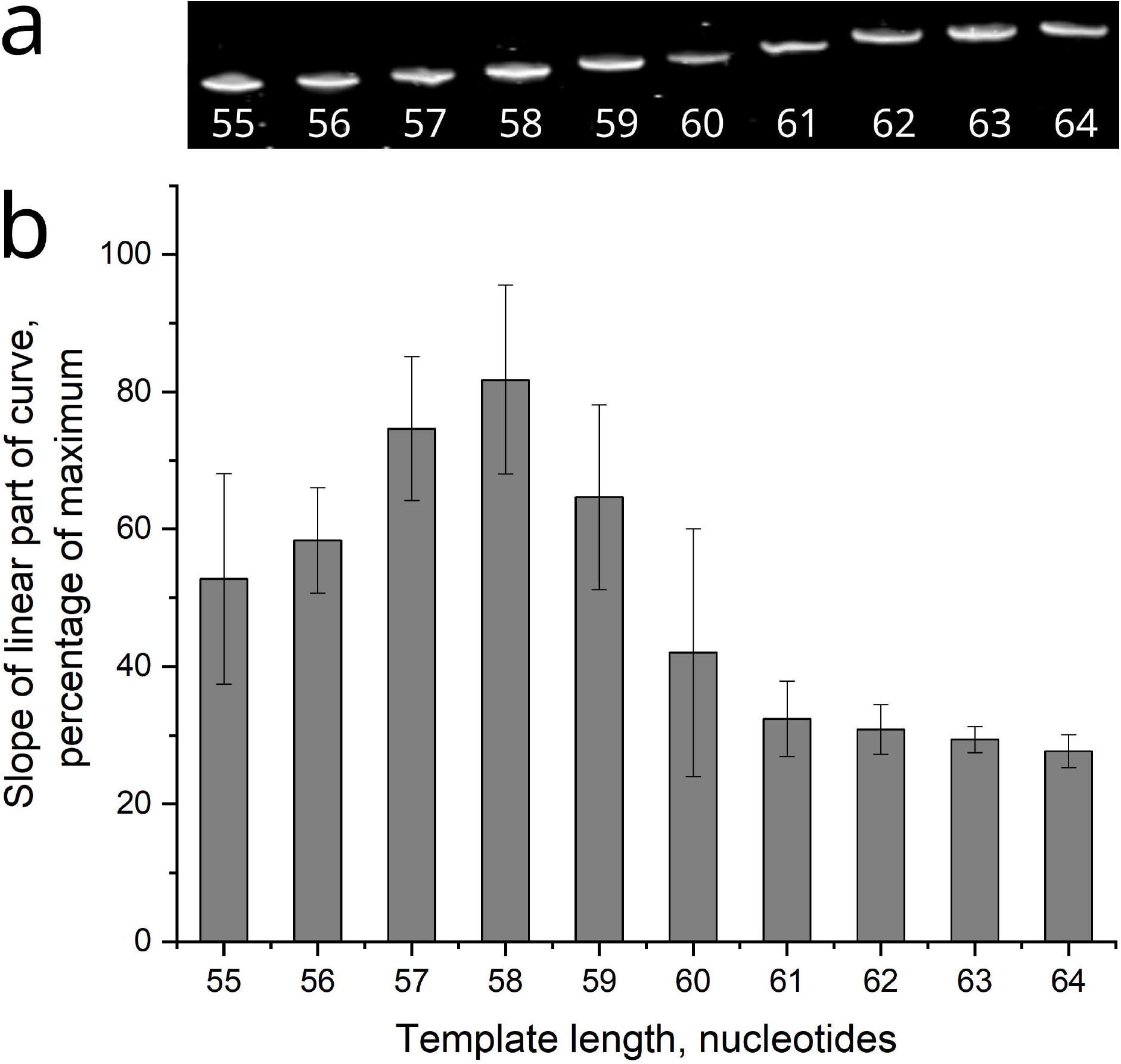

RCA maximum rate was observed with 58 pairs long circular template (Fig.1.), which is the difference between 69 bases and the length of one turn for B-DNA (10.3 bases). So, for further experiments, we took the circular template prepared from Reg58WoCt oligonucleotide.

### Immuno-RCA on the microchip

Immuno-RCA and conventional assay were compared using biotinylated anti-human antibodies – Alexa Fluor™ 555-labeled streptavidin pair (as having detection sensitivity of detection than fluorescently labeled antibodies) and human healthy donor blood group A serum as a source of detected anti-A immunoglobulins. Microchip contained A type 2 tetrasaccharide ((GalNAcα1-3(Fucα1-2)Galβ1-4GlcNAcβ) and B type 2 tetrasaccharide (Galα1-3(Fucα1-2)Galβ1-4GlcNAcβ) as a reference. The first three steps were identical both to conventional and to immuno-RCA assays (Fig. 2.) and differences in the methodology began after incubation with biotinylated secondary antibodies: in the immuno-RCA assay unlabeled streptavidin was used instead of Alexa Fluor™ 555 -labeled streptavidin followed by the application of an annealed complex of biotinylated primer with circular template. After incubation with a complex of biotinylated primer with a circular template, we carried out the RCA reaction for 1-3 hours or overnight at room temperature. During the reaction, a molecular beacon DNA probe is annealed to the single-stranded DNA molecule and begins to fluoresce. RCA product with fluorescent probes remains on the chip after washings.

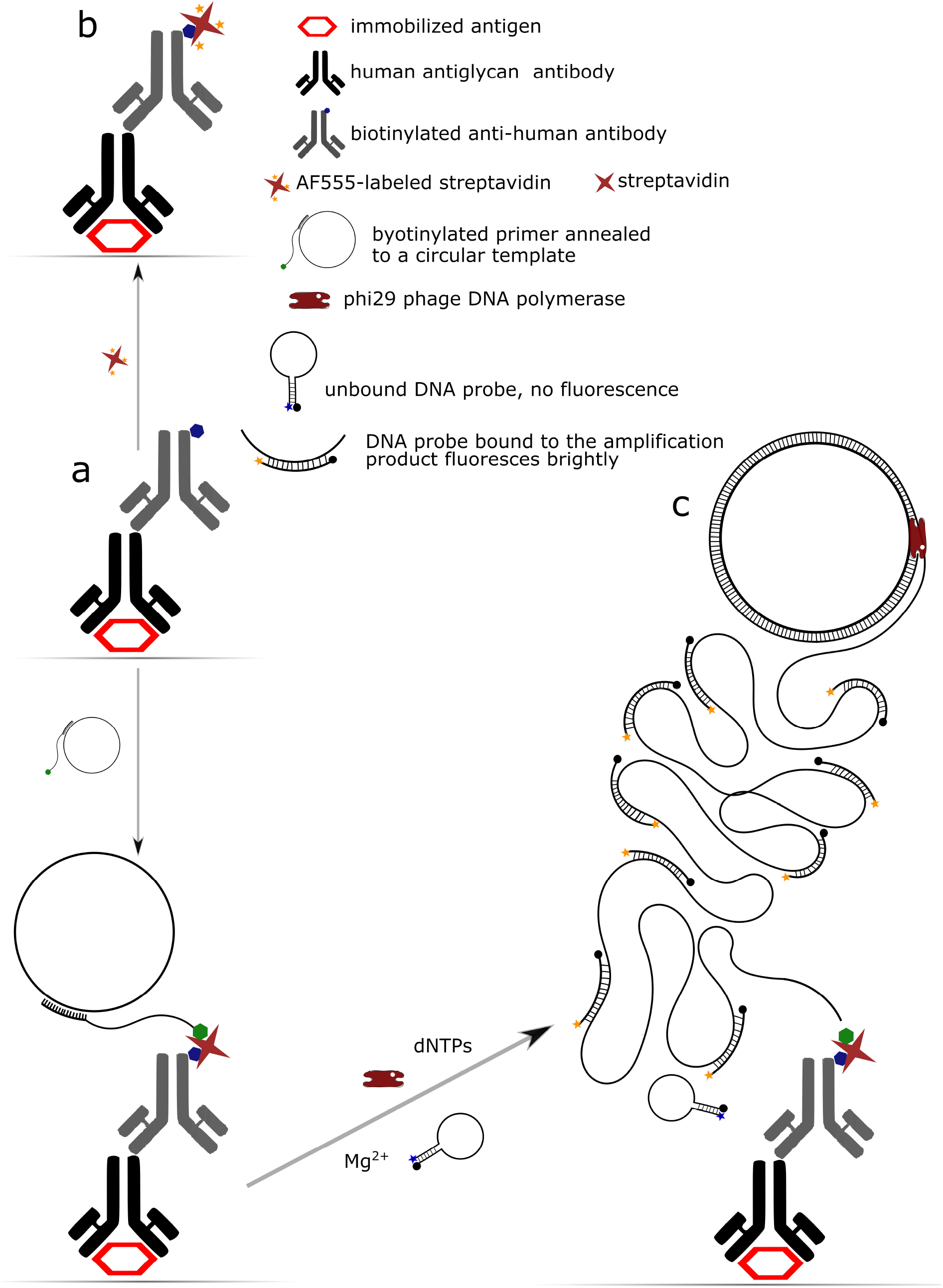

Immuno-RCA was tested with blood group antigens and natural human anti-glycan antibodies in the blood serum. The microchip consists of six printed sectors, each of them contains repeated dots with types A and B antigen, streptavidin, and BSA-Cy3 loaded in different concentrations (see Fig. S2). BSA-Cy3 and streptavidin were used to control RCA and microchip scanning equipment.

In the case of 1-hour long time RCA, were obtained the significant signals from the A type for serum dilution 1/640 while assay with Alexa Fluor™ 555 -labeled streptavidin showed positive result at 1/320 serum dilution. The greater sensitivity was achieved when the reaction was carried out overnight (Fig.3.) – at 1/5120 serum dilution. In this case, a significant (more than 3 times) ratio between fluorescence for spots with B and A antigens remains, but when the reaction mixture was mixed during the whole incubation time only.

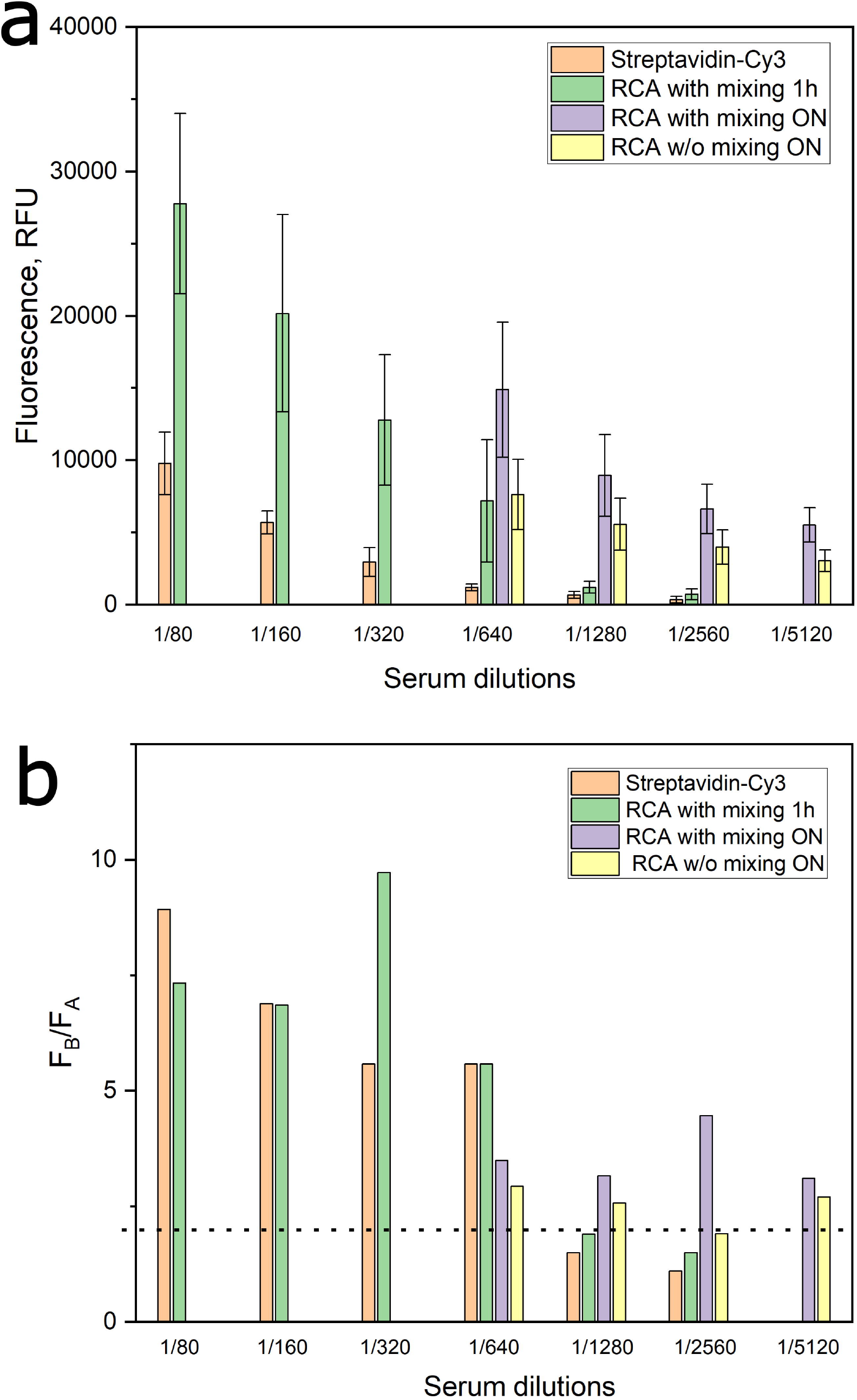

We initially expected a significant increase in fluorescence level, since RCA can produce extremely extensive amplification products that can bind hundreds of times more DNA probes than labeled streptavidin can carry. But we haven`t observed a very high fluorescence level, probably because a large part of the RCA product falls off the microchip. In the absence of mixing, the level of fluorescence and the ratio between the B-type and A-type antigens decreased, and the spots lost circular shape (see Fig. S3). The mean level of natural anti-A and anti-B antibodies (Obukhova et al., 2011) is about 5 μg/mL of serum. Thus, we achieved the anti-glycan antibody detection in the serum sensitivity of about 1 ng/mL. The key elements of successful antibody detection are the right blocking reagents (Ryazantsev et al., 2024) and concentration of the key compounds such as biotinylated antibody, streptavidin, circular template with annealed primer, RCA reaction components, duration and conditions of RCA reaction. Unfortunately, our earlier protocol (Ryazantsev et al., 2024) didn`t allow for a significant increase in antibody detection sensitivity. We were forced to increase the reaction time overnight, and thus our analysis takes about three days to increase sensitivity.

## Conclusion

We have developed a high-throughput technology for detecting anti-glycan antibodies using a printed glycan array and a rolling circle amplification reaction based on the ABO blood group antibody model. Our method significantly increased sensitivity by more than an order magnitude in contrast to conventional immunoassay with fluorescent-labeled streptavidin (from 1/320 to 1/5120 serum dilutions). However, our approach is time-consuming, as we needed to extend some incubation steps to overnight to achieve optimal results.

In the future, we will focus on evaluating the effectiveness of this technology in detecting antigen-specific antibodies and its application in identifying tumor-associated biomarkers. This innovative approach has the potential to enhance sensitivity and specificity, ultimately improving current diagnostic methods in oncology.

