## Supplemental information for "The applying of immuno-RCA for the high-sensitivity detection of the ABO blood group antibodies on the printed glycoarray"

Supplementary

Brief phage Phi29 DNA polymerase purification protocol

Gene coding phage phi29 DNA polymerase was obtained by conventional PCR gene assembly from an overlapping 60 nucleotides oligonucleotides set. The nucleotide sequence was got by the reverse translation of the P03680 amino acid sequence using the E. coli codon frequency table. The NcoI restriction enzyme site was added on the 5` end of the phi29 DNA polymerase gene sequence and 6xHis tag coding sequence followed by the HindIII restriction enzyme site on the 3` end of the phi29 DNA polymerase gene sequence. The full-length gene was digested with restriction enzymes NcoI and HindIII and ligated into pre-treated with the same enzymes pQE60 vector. Protein expression was carried out according to the following protocol. Bacteria M15 strain cells were transformed with the pQE60-p29 plasmid. The fresh LB media was inoculated by overnight bacteria culture and was grown at 42C up to 0.6 AU at 600 nm. Then, the culture was chilled up to 30C and JPTG was added up to 0.2 mM. The bacteria were cultivated after induction for about 4 h at 30C followed by the cells were collected by centrifugation. Cells were resuspended in the buffer containing 100 mM of Tris-HCl, pH 7.2, 100 mM of KCl, 0.2% of Tween-20, 1 mM of EDTA, 1 mM of PMSF, 1 mM of DTT, 1 mg/ml of lysozyme. The suspension was sonicated after incubation in ice for 30 minutes. The concentration of ammonium sulfate was adjusted up to 1M and PEI was added up to 0.1% followed by incubation in ice for 60 minutes. Clear lysate was obtained by centrifugation and applied to the column with Ni-NTA resin (Qiagen). The column was washed with 2 volumes of starting buffer (100 mM Tris-HCl, pH 7,2, 100 mM KCl, 0.05% Tween-20, 1 М ammonium sulfate, 20 mM imidazole) and wash buffer (100 mM Tris-HCl, pH 7,2, 100 mM KCl, 0.05% Tween-20, 1 М ammonium sulfate, 10 mM imidazole) consequentially. The polymerase was eluted with MAC elution buffer (100 mM Tris-HCl, pH 7,2, 100 mM KCl, 0.05% Tween-20, 1 М ammonium sulfate, 250 mM imidazole) and applied to the column with phenyl-sepharose CL-4B resin (Amersham). The column was washed with 2 volumes of wash buffer (100 mM Tris-HCl, pH 7,2, 100 mM KCl, 0.05% Tween-20, 0.5 М ammonium sulfate) and the polymerase was eluted with HC elution buffer (100 mM Tris-HCl, pH 7,2, 100 mM KCl, 0.05% Tween-20, 0.2 М ammonium sulfate) and concentrated on Corning Spin-X concentrator (30 kDa). Concentrated lysate was applied to Hiload Superdex 200 16/600 pg column (Cytiva) equilibrated with buffer containing 100 mM Tris-HCl, pH 7,2, 100 mM KCl, 0.2% Tween-20. Eluate was supplemented with  0.2% NP-40,1 mM DTT, concentration was measured and glycerol was added up to 50%.

Table S1. List of oligonucleotides used

| # | Oligonucleotide | Sequence, 5`-3` |
| --- | --- | --- |
| 1 | lockWoC | GATAAATGGGTAAATA[Alkyne] |
| 2 | Reg64WoCt | [p]CATTTATCTATTCACCTATTTATTCACCCACTTACTTACAACCACACCCACTCACTTATTTACC |
| 3 | Reg63WoCt | [p]CATTTATCTATTCACCTATTTATTCACCCACTTACTTACAACACACCCACTCACTTATTTACC |
| 4 | Reg62WoCt | [p]CATTTATCTATTCACCTATTTATTCACCCACTTACTTACACACACCCACTCACTTATTTACC |
| 5 | Reg61WoCt | [p]CATTTATCTATTCACCTATTTATTCACCCACTTACTTACACCACCCACTCACTTATTTACC |
| 6 | Reg60WoCt | [p]CATTTATCTATTCACCTATTTATTCACCCACTTACTTACCCACCCACTCACTTATTTACC |
| 7 | Reg59WoCt | [p]CATTTATCTATTCACCTATTTATTCACCCACTTACTTACCACCCACTCACTTATTTACC |
| 8 | Reg58WoCt | [p]CATTTATCTATTCACCTATTTATTCACCCACTTACTTACACCCACTCACTTATTTACC |
| 9 | Reg57WoCt | [p]CATTTATCTATTCACCTATTTATTCACCCACTTACTTCACCCACTCACTTATTTACC |
| 10 | Reg56WoCt | [p] CATTTATCTATTCACCTATTTATTCACCCA CTTACCAC CCACTCACTTATTTACC |
| 11 | Reg55WoCt | [p]CATTTATCTATTCACCTATTTATTCACCCA CTTACAC CCACTCACTTATTTACC |
| 12 | pr25WoCtBio | [bio]TTTTTTTTTTTTTTTTTTTGATAAATGGGTAAATAAGTGAGTGG |
| 13 | beacWoC3 | [Cy3]CCGGGTATCTATTCACCTATTTATTCACCCGG[BHQ2] |

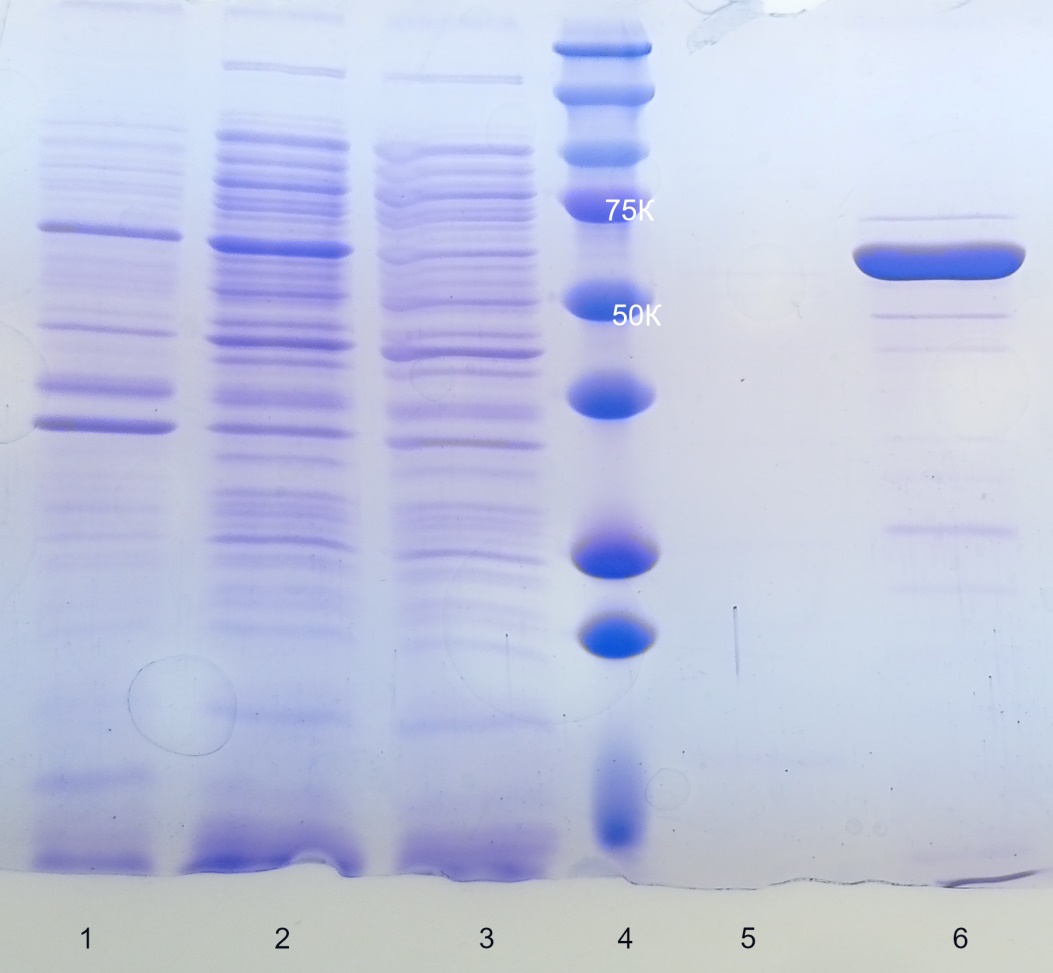

Fig. S1.. SDS-PAGE of recombinant phi29 phage DNA polymerase fractions

1. Cells lysate before induction, 2- cells lysate after induction, 3- Ni-NTA column flow-through, 4 - Precision Plus Protein Standards (Bio Rad), 5 - Ni-NTA column wash fraction, 6 - Ni-NTA column elution fraction.

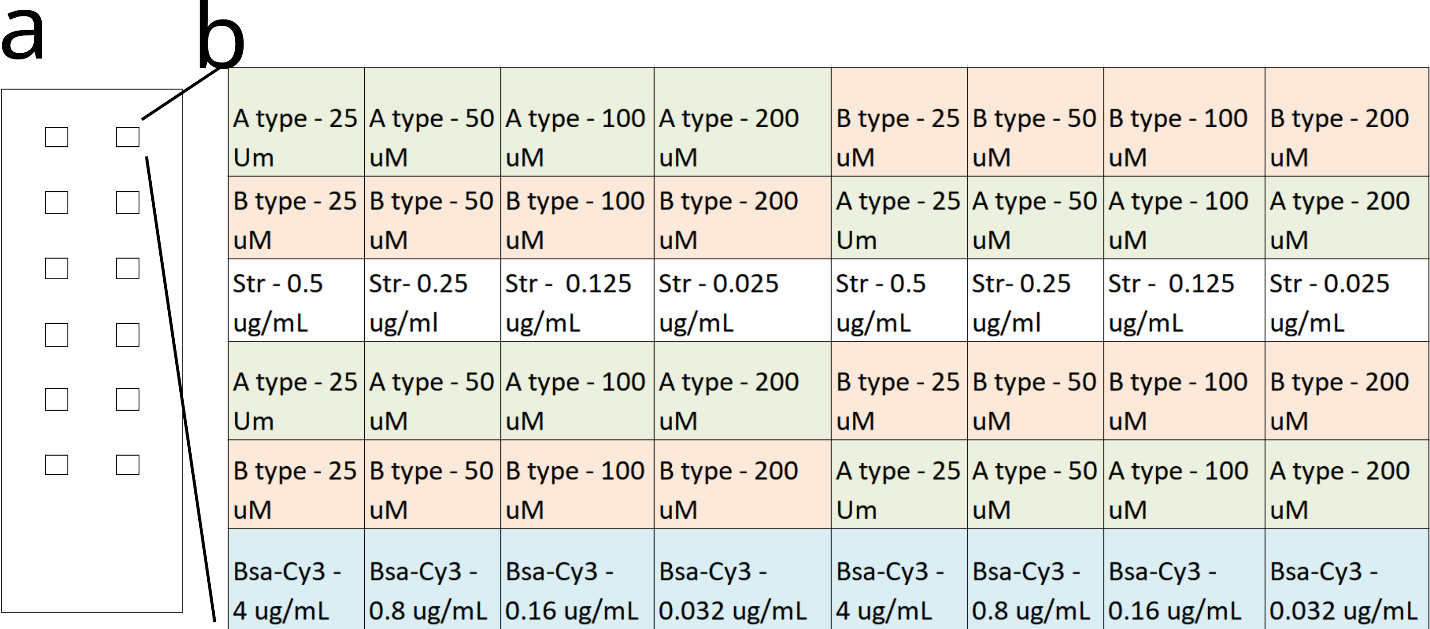

Fig. S2. Scheme of the whole microchip (a) and separate printing sector on it (b).

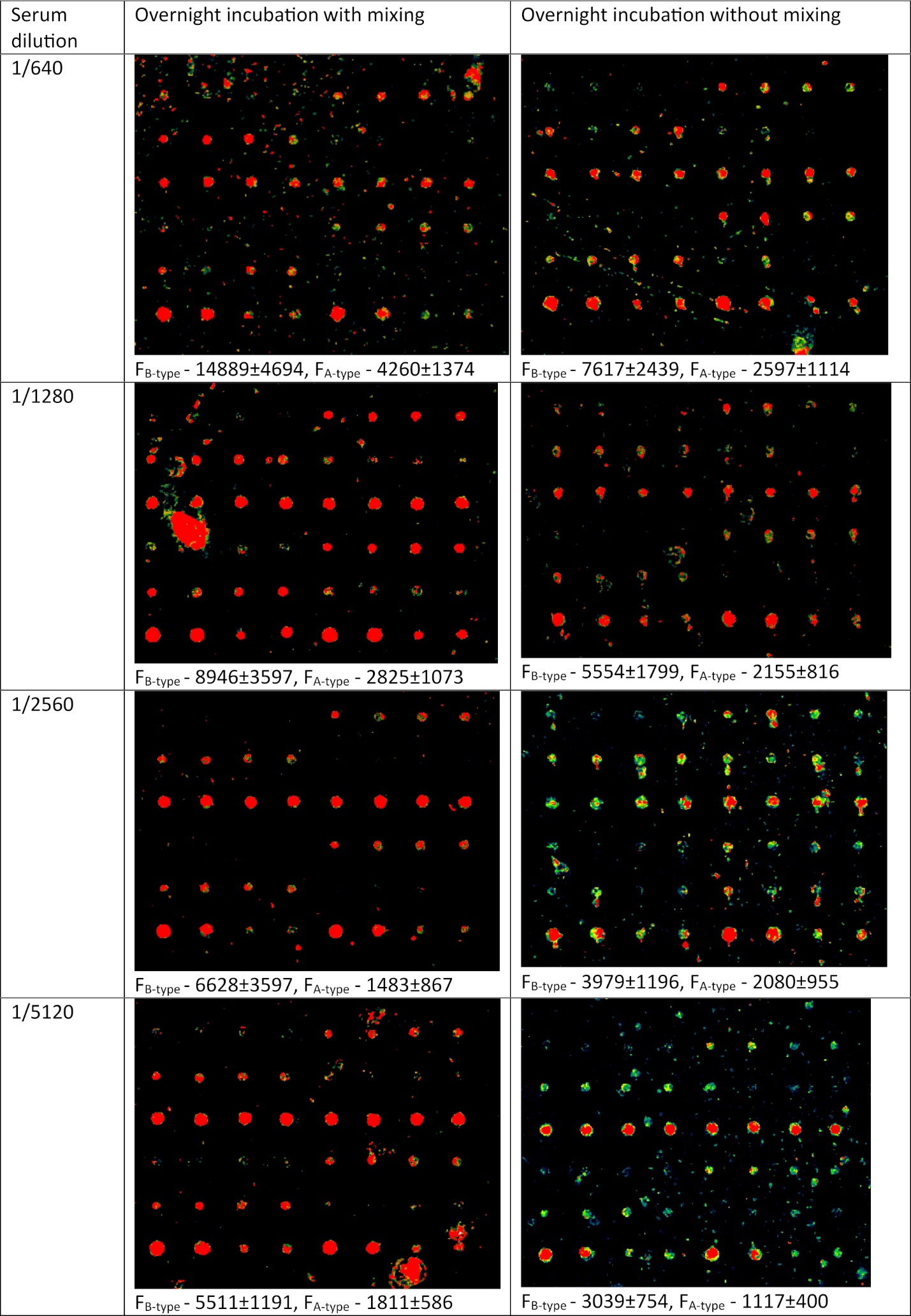

Fig.S3. Typical segments of microchip with different RCA protocols.

Phi29 phage DNA polymerase gene sequence used

**ccatgg**atgaaacatatgccgcgtaagatgtatagctgcgatttcgaaaccaccaccaaagtggaagattgccgtgtgtgggcatatggctatatgaacattgaagatcatagcgaatataagattggcaacagcctggatgaatttatggcgtgggtgctgaaggtgcaggcagacctgtatttccataacctgaagtttgatggcgcattcattatcaactggctggaacgtaacggcttcaagtggagcgcagatggtctgccgaacacctataacaccattatcagccgtatgggtcagtggtatatgattgatatttgcctgggctataaaggcaaacgtaagattcataccgtgatctatgatagcctgaagaaactgccgtttccggtgaagaagattgcgaaggatttcaaactgaccgtgctgaaaggcgatatcgattatcataaggaacgtccggtgggttataagatcaccccggaagaatatgcgtatatcaagaacgatattcagatcatcgcagaagcactgctgattcagttcaaacagggcctggatcgtatgaccgcaggtagcgatagcctgaaaggcttcaaggatatcatcaccaccaagaagttcaagaaggtgtttccgaccctgagcctgggactggataaggaagtgcgttatgcatatcgtggtggtttcacctggctgaacgatcgtttcaaggaaaaggaaattggcgaaggcatggtgtttgatgtgaacagcctgtatccggcgcagatgtatagccgtctgctgccgtatggcgaaccgattgtgttcgaaggcaaatatgtgtgggatgaagattatccgctgcatattcagcatattcgttgcgaatttgaactgaaagaaggctatattccgaccattcagatcaaacgtagccgtttctataagggcaacgaatatctgaaaagcagcggaggcgaaattgcggatctgtggctgagcaacgtggatctggaactgatgaaagaacattatgatctgtataacgtggaatatattagcggcctgaagttcaaagcgaccaccggactgttcaaggatttcattgataagtggacctatatcaagaccaccagcgaaggtgcgatcaagcagctggcgaaactgatgctgaacagcctgtatggcaagtttgcgagcaacccggatgtgaccggcaaagtgccgtatctgaaggaaaacggcgcactgggctttcgtctgggtgaagaagaaaccaaagatccggtgtataccccgatgggtgtgttcattaccgcatgggcacgttataccaccattaccgcagcacaggcatgctatgatcgtatcatctattgcgataccgatagcatccatctgaccggcaccgaaatcccggatgtgatcaaggatatcgtggacccgaagaagctgggctattgggcacatgaaagcaccttcaagcgtgcgaagtatctgcgtcagaagacctatatccaggatatctatatgaaagaagtggatggcaagctggtggaaggtagcccggatgattataccgatatcaagttcagcgtgaaatgcgcgggtatgaccgataaaatcaagaaggaagtgaccttcgaaaacttcaaggtgggctttagccgtaagatgaaaccgaaaccggtgcaggtgccgggtggtgtggtgctggtggatgataccttcaccatcaagagatctcatcaccatcaccatcact**aagctt**
